# From a deep learning model back to the brain - inferring morphological markers and their relation to aging

**DOI:** 10.1101/803742

**Authors:** Gidon Levakov, Gideon Rosenthal, Ilan Shelef, Tammy Riklin Raviv, Galia Avidan

**Author notes:** Equal contribution (Tammy Riklin Raviv, Galia Avidan). Data used in preparation of this article was partially obtained from the Alzheimer’s Disease Neuroimaging Initiative (ADNI) database (adni.loni.usc.edu) and the Australian Imaging Biomarkers and Lifestyle flagship study of ageing (AIBL) funded by the Commonwealth Scientific and Industrial Research Organisation (CSIRO). As such, the investigators within ADNI and AIBL contributed to the design and implementation of ADNI and AIBL and/or provided data but did not participate in analysis or writing of this report. A complete listing of ADNI and AIBL investigators can be found at http://adni.loni.usc.edu/wp-content/uploads/how_to_apply/ADNI_Acknowledgement_List.pdf and at www.aibl.csiro.au. Corresponding author. E-mail address (G. Levakov).

## Abstract

We present a Deep Learning framework for the prediction of chronological age from structural MRI scans. Previous findings associate an overestimation of brain age with neurodegenerative diseases and higher mortality rates. However, the importance of brain age prediction goes beyond serving as biomarkers for neurological disorders. Specifically, utilizing convolutional neural network (CNN) analysis to identify brain regions contributing to the prediction can shed light on the complex multivariate process of brain aging. Previous work examined methods to attribute pixel/voxel-wise contributions to the prediction in a single image, resulting in ‘explanation maps’ that were found noisy and unreliable. To address this problem, we developed an inference framework for combining these maps across subjects, thus creating a population-based rather than a subject-specific map. We applied this method to a CNN ensemble trained on predicting subjects’ age from raw T1 brain images of 10,176 subjects. Evaluating the model on an untouched test set resulted in mean absolute error of 3.07 years and a correlation between chronological and predicted age of r=0.98. Using the inference method, we revealed that cavities containing CSF, previously found as general atrophy markers, had the highest contribution for age prediction. Comparing maps derived from different models within the ensemble allowed to assess differences and similarities in brain regions utilized by the model. We showed that this method substantially increased the replicability of explanation maps, converged with results from voxel-based morphometry age studies and highlighted brain regions whose volumetric variability contributed the most to the prediction.

**Highlights:**

- CNNs ensemble is shown to estimate “brain age” from sMRI with an MAE of ∼3.1 years
- A novel framework enables to highlight brain regions contributing to the prediction
- This framework results in explanation maps showing consistency with the literature
- As sample size increases, these maps show higher inter-sample replicability
- CSF cavities reflecting general atrophy were found as a prominent aging biomarker

## 1. Introduction

The human brain undergoes complex structural changes across the lifespan (Sowell, Thompson, & Toga, 2004). These include widespread synaptic pruning and myelination from early life through puberty and neurodegenerative processes, such as ventricle expansion and cortical thinning that peaks at aging. The course and extent of these changes are not uniformly distributed across the brain (Storsve et al., 2014). Thus, for example in healthy aging, higher atrophy rates were reported in the hippocampus, while regions like the early visual cortex remain relatively intact (but see: Lemaitre et al., 2012). Nevertheless, studies that examined the correspondence between brain structure and chronological age provide inconsistent findings. Such inconsistencies may be related to the specific parcellation schemes employed (Mikhael & Pernet, 2019), surface-based structural measurements (Lemaitre et al., 2012) or global volume covariates (Jäncke, Mérillat, Liem, & Hänggi, 2015). These concerns add to discrepancies due to the usage of relatively small samples and different statistical procedures, which together impede the attempts to characterize the relation between aging and structural changes in the brain.

Studying brain aging has important implications for differentiating typical and pathological aging. Alzheimer’s disease (AD), the most prevalent type of dementia, affects about 22% of the population over the age of 75 (in the US, 2010; Hebert, Weuve, Scherr, & Evans, 2013). AD patients exhibit extensive cell loss in cortical and subcortical regions, but such findings are also evident in typical aging (Arendt, Brückner, Morawski, Jäger, & Gertz, 2015). Moreover, behavioral manifestations such as cognitive decline and memory deficits that accompany AD, are also apparent in aging in the absence of AD (Cardenas et al., 2011; Koen & Yonelinas, 2014). Thus, a reliable measure of typical brain aging may be beneficial in order to better distinguish between the two (Lorenzi, Pennec, Frisoni, & Ayache, 2015).

### 1.1. Predicting age from structural brain imaging using machine learning

Recent growth in data availability and advancements in the field of machine learning (ML), applied to the analysis of structural imaging, have allowed addressing regression problems such as brain age prediction based on preselected sets of anatomical features or regions of interest (ROIs). Predicting age from brain anatomy enables to estimate a measure of “brain age” which is independent of one’s chronological age. Different studies generally reveal that an over-estimation of that measure is associated with neurodegenerative diseases and various clinical conditions and might even predict mortality (Cole et al., 2018). Hence, brain age estimation could be used as a potential biomarker for brain health (Cole & Franke, 2017). While ML methods were shown to provide a mean error of ∼5 years (Cole & Franke, 2017), age predictions are largely dependent upon the selection of features that would be used as input to the algorithm.

### 1.2. Application of deep Convolutional neural network for predicting “brain age”

Deep Convolutional neural network (CNN) has enabled a major leap in many applications including neuroimaging analysis, among others, by learning the features, or representation from the raw data, i.e., an image or a volume (Goodfellow, Bengio, & Courville, 2016). CNNs are biologically inspired algorithms in which the connectivity between the different neurons implements a convolution operation. The neurons are ordered in stacked layers in a hierarchical deep formation and hence they are termed deep CNN (LeCun, Bottou, Bengio, & Haffner, 1998). CNN based models achieved state-of-the-art results in serval neuroimaging tasks including cortical segmentation and tumor detection (Kamnitsas, Chen, & Ledig, 2015; Pereira, Pinto, Alves, & Silva, 2016) and were recently applied to age prediction from raw T1 magnetic resonance images (MRI) images (Cole, Poudel, et al., 2017). Nonetheless, significant improvement can still be achieved by substantially increasing the sample size and utilizing practices such as prediction based on an ensemble of models. Both of these approaches were shown to produce a remarkable improvement in other visual task domains (Lee, Purushwalkam, Cogswell, Crandall, & Batra, 2015).

### 1.3. Model interpretability – which brain regions underlie a given prediction?

A major limitation of studies utilizing CNNs, pertains to the issue of the model interpretability. While CNNs have provided high accuracy for age prediction (Cole & Franke, 2017; Qi, Du, Zhuang, Huang, & Ding, 2018), it is typically difficult to identify the features that enabled a given prediction. Several recent studies attempted to identify or visualize intermediate representations of the CNN (Olah et al., 2018), but still, the size and complexity of the networks make it a challenging task. In the context of structural neuroimaging analysis, there might be an advantage to focus on the input level since it could be directly related to specific brain structures. Knowing which image parts, or in the current research, brain regions or neural attributes, contribute most to the prediction have theoretical as well as translational value. A possible approach to this issue is the usage of “saliency maps” or “explanation maps” indicating the influence of each voxel in the input volume on the model’s prediction. Such a map can be generated by calculating the partial derivative of each voxel in the input volume with respect to the model’s output (Simonyan, Vedaldi, & Zisserman, 2013; Springenberg, Dosovitskiy, Brox, & Riedmiller, 2014). However, local gradients in non-linear functions such as CNN were previously shown to be noisy. Recent work has demonstrated that this could be partially addressed by repeatedly calculating and averaging several explanation maps derived from the same input after adding random noise to it (Smilkov, Thorat, Kim, Viégas, & Wattenberg, 2017). Nevertheless, these explanation maps are typically created on a single sample, hence they provide only a subject-specific rather than a population-based explanation (Yang, Rangarajan, & Ranka, 2018; but see: Bermudez et al., 2019; Wang et al., 2019). In a task or a model were large variability exists in these explanation maps, i.e., if different subject-level maps highlight different regions, any translational or theoretical conclusion drawn from it could only be subject-specific.

### 1.4. The current study

In light of the limitations outlined above, we aimed to examine brain aging using a CNN model for “brain age” prediction and identify the brain structures that supported this prediction. Therefore, this study has two important contributions. The first is the prediction model, which is composed of an ensemble of multiple CNNs trained to predict individuals’ age from minimally processed T1 MRI scans. The model was trained and tested on an aggregated sample size of 10,176 subjects, from several large-scale open-access databases (n = 15), producing a result robust to scanner’s type, field strength, and resolution. Second, we provided and validated a novel framework for identifying the importance of the various anatomical brain regions to the prediction by aggregating multiple subject-level explanation maps, creating a population-based map. Combining subject-level maps into a population-based map is done by image realignment after training the model, thus no special preprocessing or architecture modification is required, as opposed to previous work (Ito et al., 2018). We empirically show that this significantly improves the explanation maps and allows the inference from the model back to the brain’s anatomy. The use of an ensemble of CNNs, apart from an increase in prediction accuracy, allows examining the diversity and the similarity of independently trained models, or to what extent different models exploit similar brain regions for the age prediction.

## 2. Material and methods

### 2.1. Datasets

To train a model that is robust to different sites and scanning protocols, we collected a dataset of T1w MRI brain scans of 10,176 individuals from various open-databases (n = 15), acquired at different locations, scanners, and scanning parameters. Several databases from longitudinal studies consist of brain scans acquired at several time-points. For these databases, we only used scans of the first time point to avoid data leakage between the train and validation/test sets. Three exclusion criterions were applied to all subjects: missing age report, major artifacts in a visual inspection of the T1 volume and diagnosis of AD or another form of dementia. The complete list of studies, age, and gender distributions are reported in Table 1.

**Table 1.** List of all studies which comprise the dataset. For each study, the number of available subjects (N), the mean and standard deviation of the age and gender distribution are provided. * To prevent participant identification in the GSP study age was rounded to the closest 2 years bin.

| **<u>Study/database</u>** | **<u>N</u>** | **<u>Age M(± SD)</u>** | **<u>Gender (F:M)</u>** |
| --- | --- | --- | --- |
| Consortium for Reliability and Reproducibility (CoRR; Zuo et al., 2014) | 1378 | 26.0 (±15.8) | 693; 685 |
| Alzheimer's Disease Neuroimaging Initiative (ADNI; Jack et al., 2008) | 1476 | 73.0 (±7.0) | 563; 912 |
| Brain Genomics Superstruct Project (GSP; Buckner, Roffman, & Smoller, 2014)* | 1099 | 21.5 (±2.9) | 630; 469 |
| Functional Connectomes Project (FCP; Biswal et al., 2010) | 1067 | 28.9 (±13.9) | 594; 473 |
| Autism Brain Imaging Data Exchange (ABIDE; Di Martino et al., 2014) | 1053 | 17.1 (±8.1) | 153; 900 |
| Parkinson's Progression Markers Initiative (PPMI; Marek et al., 2011) | 702 | 61.7 (±10.2) | 260; 442 |
| International Consortium for Brain Mapping (ICBM; Mazziotta, Toga, Evans, Fox, & Lancaster, 1995) | 641 | 30.6 (±12.2) | 293; 348 |
| Australian Imaging, Biomarkers and Lifestyle (AIBL; Ellis et al., 2009) | 616 | 72.9 (±6.6) | 342; 273 |
| Southwest University Longitudinal Imaging Multimodal (SLIM; Liu, Wei, Chen, Yang, & Meng, 2017) | 574 | 20.1 (±1.3) | 320; 252 |
| Information extraction from Images (IXI; Heckemann et al., 2003) | 563 | 48.2 (±16.5) | 312; 252 |
| Open Access Series of Imaging Studies (OASIS; Marcus, Fotenos, Csernansky, Morris, & Buckner, 2010; Marcus et al., 2007) | 402 | 51.6 (±24.9) | 257; 145 |
| Consortium for Neuropsychiatric Phenomics (CNP; Poldrack et al., 2016) | 252 | 33.3 (±9.3) | 112; 153 |
| Center for Biomedical Research Excellence (COBRE; Mayer et al., 2013) | 146 | 37.0 (±12.8) | 37; 109 |
| Child and Adolescent NeuroDevelopment Initiative (CANDI; Frazier et al., 2008) | 103 | 10.8 (±3.1) | 46; 57 |
| Brainomics (Pinel et al., 2012) | 89 | 24.7 (±6.8) | 47; 42 |
| **Overall** | **10174** | **39.4 (±23.8)** | **4659; 5511** |

### 2.2. Data preprocessing

To minimize the model reliance on preprocessing steps such as image realignment and registration that are both computationally intensive and time-consuming, we designed a minimal preprocessing procedure. To ensure that the model “brain age” estimation would rely solely on regions within the skull, the only substantial preprocessing step was the removal of extra-cranial regions from the volume. Thus, the preprocessing procedure included 4 stages: Applying a coarse (90°) rotation so that all the volumes would appear in similar L-R, A-P, S-I orientation (FSL *fslreorient2std* tool; Woolrich et al., 2009), skull removing tool (ROBEX; Iglesias, Cheng-Yi Liu, Thompson, & Zhuowen Tu, 2011), volume resize to standard size (90, 126, 110) and volume standardization (μ=0, σ=1). Resizing of each volume was implemented by applying an identical scaling factor to all 3 dimensions, such that brain voxels (> 0) would occupy the maximum portion within the final volume (90, 126, 110). For each volume, voxels’ intensities were standardized by a subtraction of the volume’s mean intensity followed by division by the intensity’s standard deviation.

### 2.3. Data augmentation

Head orientation, the field of view (FOV) and the level of signal to noise ratio (SNR) may differ between scans even if they were acquired by the same machine and are of the same subject. To improve the robustness of the models to these variations we augmented the training data by randomly manipulating the head position, size, and noise level. This procedure was previously shown to improve generalization and avoid overfitting (Simard, Steinkraus, & Platt, 2003). Specifically, the series of transformation to the brain image included rotation in the x/y/z-axis *unif*(−10°, 10°), shifting *unif*(−5,5) voxels, scaling 𝒩(0,0.1) and adding random noise 𝒩(0,0.015). The optimal augmentation parameters were chosen as the ones that maximized the validation accuracy using a random hyperparameter search.

### 2.4. CNN architecture

The CNN models were implemented using Keras (François Chollet and contributors, 2015) with TensorFlow (Abadi et al., 2016) backend. Each 3D CNN model was trained separately to predict age from a T1 MRI. The input for each network was a 3D volume, of size [90, 126, 110] and the output was a single scalar representing chronological age (years). The model was composed of 2 blocks, each with a batch normalization layer (Ioffe & Szegedy, 2015) followed by two 3D convolutional layers and a max-pooling layer. The two blocks were followed by 2 fully connected layers (FC). All layers, but the last fully connected one were followed by a ReLU non-linear activation (Nair & Hinton, 2010). To reduce overfitting, we added dropout layers after the convolutional layer and before the last layer for the training stage (see Fig. 1a for the complete architecture). The loss function for each CNN was the mean squared error between the real and predicted age. The network architecture was chosen using random hyperparameters search.

**Figure 1.**
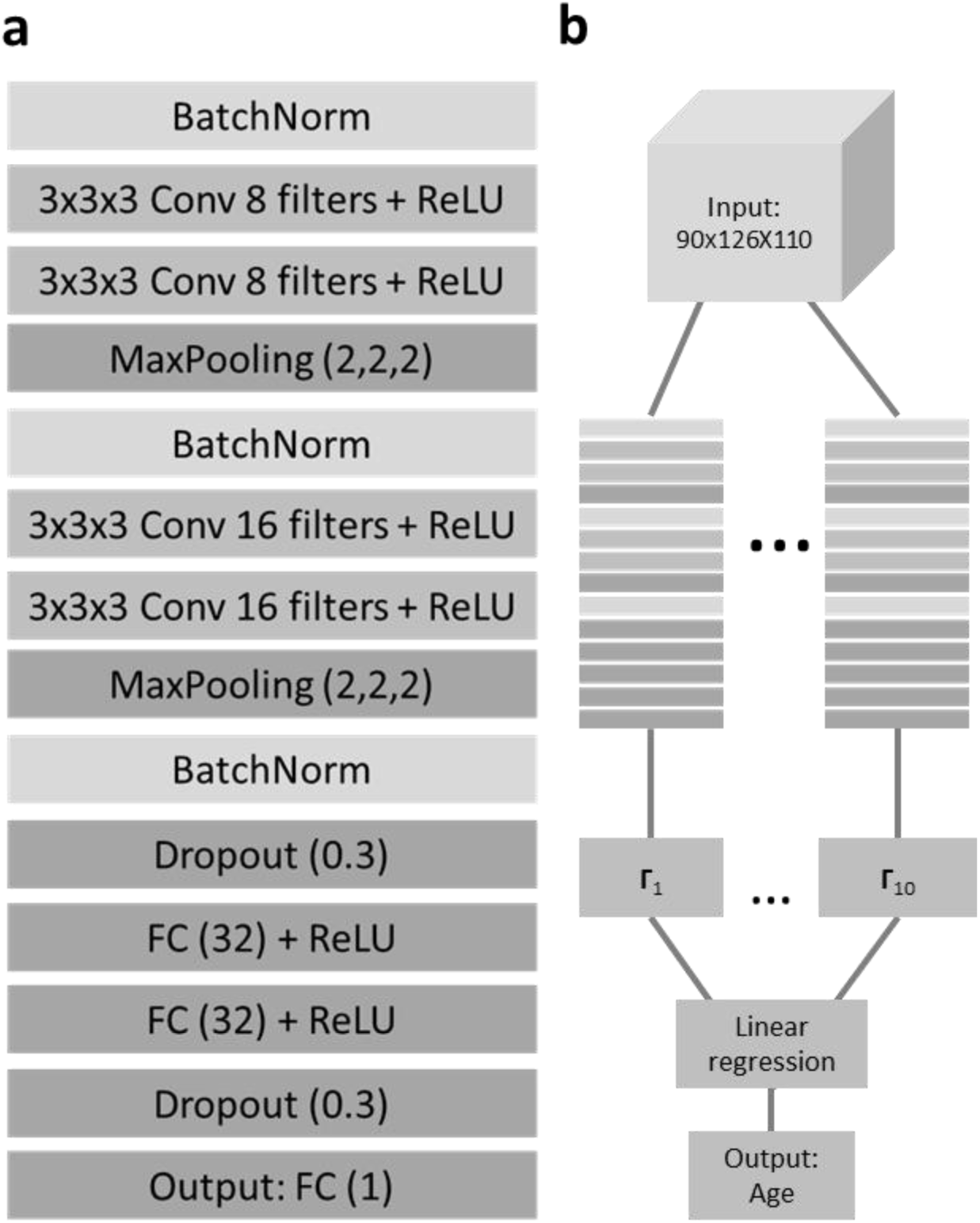
Network architecture for age prediction. (a) The detailed architecture of the network used for age prediction from 3D T1 MRI volume. BatchNorm = batch normalization, Conv = convolutional layer, = rectified linear unit, FC = fully connected layer. (b) The ensemble procedure combining the output of ten separately trained CNNs (Γ_1-10_) using linear regression to create the final age prediction.

### 2.5. The ensemble model

The ensemble in the current work included multiple 3D CNNs (*m* = 10) each trained separately to predict age from a T1 MRI. As in previous work utilizing CNNs (Lakshminarayanan, Pritzel, & Blundell, 2017; Lee et al., 2015), ensemble models differ only in their random weight initialization. Hence, they had identical architecture and were trained on the same data set. We picked m = 10 due to consideration of training time given the large number of parameters in a 3D CNN and the large training set. After each network was independently trained, a linear regression model for age prediction is learned from the outputs of the ten networks using the same training set (see Fig. 1b). In Section 2.9 we describe the measures of the models’ similarity/diversity using our proposed population-based explanation map.

### 2.6. Performance metrics

All databases were randomly divided into training (90%), validation (5%) and test (5%) sets. The training set was used to train each network separately and to find the optimal ensemble weights. The validation set was used for hyperparameters tuning and to assess over-fitting. All performance measures were calculated on the untouched test set. The prediction was evaluated using mean absolute error (MAE) and the Pearson correlation coefficient between the network prediction and the chronological age values.

### 2.7. Individual explanation maps

We employed the SmoothGrad method (Smilkov et al., 2017) that was implemented using iNNvestigate (Alber et al., 2018). This is a gradient-based method in which a given input image is first distorted with random noise from a normal distribution *N*(μ=0, σ=0.1), then the partial derivative of each voxel is computed with respect to the trained model’s output. This was repeated several times (k = 32), then the produced gradient maps were averaged. We used partial derivative following Adebayo et al., (2018) work that demonstrated that it best captures the CNN’s training process.

### 2.8. Aggregating explanation maps across samples

First, the models preprocessed input was transformed into the raw anatomical space using FSL FLIRT (Jenkinson, Bannister, Brady, & Smith, 2002) followed by surface-based non-linear registration to the MNI space using Freesurfer (Greve & Fischl, 2009). The transformations were computed on the T1 images, then applied to the explanation maps. The complete pipeline was created using Nipype (Gorgolewski et al., 2011). Next, each volume is standardized (μ=0, σ=1), and smoothed with a 3D Gaussian using Scikit-image (Full width at half maximum = 4; van der Walt et al., 2014). Finally, all volumes were averaged to create population-based explanation maps. In line with previous work, we used the absolute value of the resulting maps (Ancona, Ceolini, Öztireli, & Gross, 2017). To identify regions with the highest contribution to the model’s prediction, we threshold the map, keeping only 1% of the voxels with the highest gradient value (see fig. 2 – for the inference scheme). To create an ensemble population-based map, we aggregate these population-based maps generated for each of the ten CNN by taking the median of each voxel across the ten maps. We will refer to the statistics obtained for these explanation maps, which is the standardized partial derivative, as an *explanation score* (ES).

**Figure 2.**
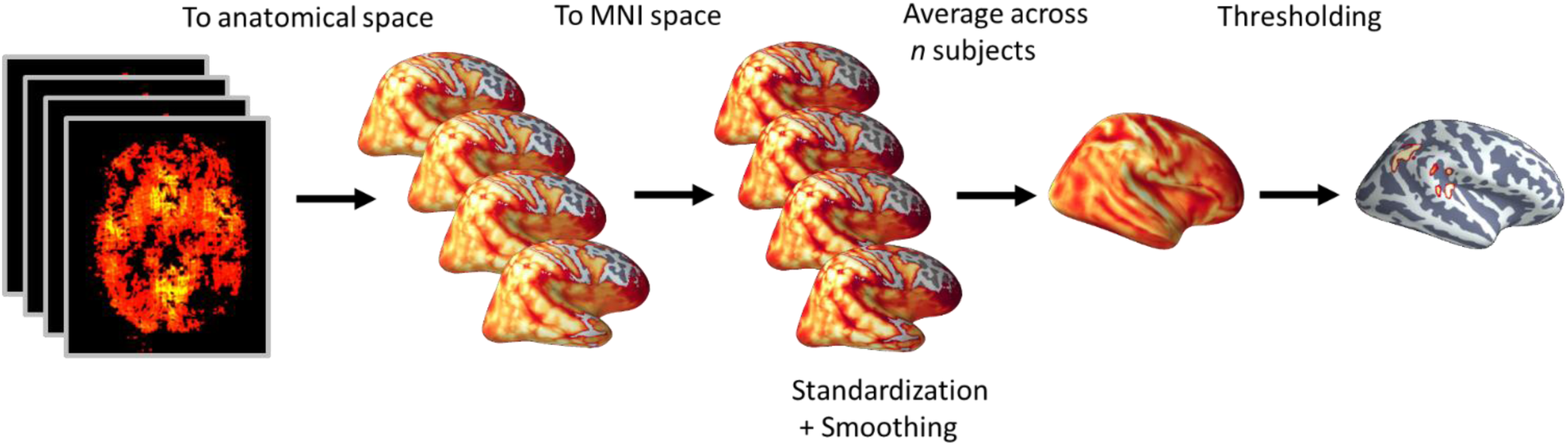
A layout of the inference scheme. For a subset of *n* subjects, an explanation map was computed, representing the contribution of each voxel to the model’s output. Each saliency map was first registered to the subject anatomical image, then it was transformed to the MNI space. Next, each volume was smoothed with a 3D Gaussian. Finally, all the volumes were averaged to create a population-based explanation map.

### 2.9. Assessing the similarity of explanation maps within the ensemble

To assess the diversity among independently trained CNNs, or the extent to which different CNNs utilize different brain regions for the prediction, we examined the similarity among their explanation maps. Specifically, the similarity between each pair of population-based explanation maps (n = 100) was evaluated with two measures: Dice similarity (Zou et al., 2004) and the modified Hausdorff distance (MHD; Dubuisson & Jain, 2002) on the threshold maps. Maps thresholding was generated by taking the absolute value of each population-based map, computing the 5^th^ percentile of the ES within the brain mask and creating a binarized map for super-threshold values: 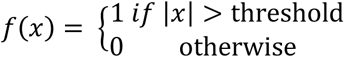. For each pair of binarized maps, the Dice coefficient was defined as: 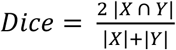, where |*X* ∩ *Y*| is the number of overlapping super-threshold voxels in both maps, and |X| and |Y| are the number of super-threshold voxels for maps X and Y, respectively. MHD was derived by first finding the surface for each cluster of super-threshold voxels within each map by using a gradient-based edge detector. Then, the MHD, or the symmetric average surface distance was calculated as follows:

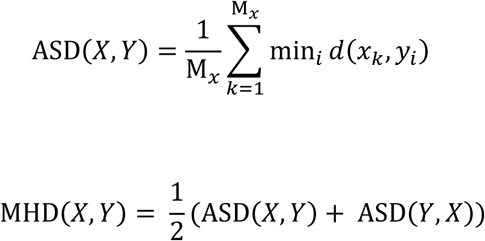

Where *d* is the Euclidian distance, M_x_ is the number of voxels in the surface map X and x and y are points on the surface in maps X and Y, respectively. Both the MHD and Dice coefficients were calculated for each pair of maps creating a distance/similarity matrix. The mean distance/similarity was calculated by taking the mean over the lower triangle of that matrix.

### 2.10. Relating contribution to specific tissues and brain structures

To obtain a general view of the features utilized by the model, we first segmented the brain volume to 4 classes of tissue type: cerebrospinal fluid (CSF) and choroid plexus, white matter (WM), subcortical gray matter (GM) and cortical GM. These were determined by applying the Desikan-Killiany Atlas (Desikan et al., 2006) using Freesurfer on the MNI template. Taking the calculated ensemble population-based explanation map, we devised a volume-normalized class ES by dividing the mean ES in each class by the total class volume. The resulting class scores were reported as a percentage of the sum of class scores. Next, to identify the specific brain regions that contributed the most for predicting age, we identified clusters of voxels in the threshold map (1^st^ percentile) using FSL cluster (Woolrich et al., 2009). For each cluster, we report the name of the brain region, its MNI coordinates, the cluster size and peak ES within the cluster. Brain regions were identified by locating the pick value of each cluster in the Desikan-Killiany Atlas for GM structures and with the ICBM-DTI-81 Atlas (Mori, Wakana, Zijl, & Nagae-Poetscher, 2005) for WM structures. Since many of the clusters were located within CSF spaces, whose sub-parts are poorly delineated in most parcellation, we manually identified sub-divisions of the cisterns and ventricles. The unthresholded population-based maps for each of the ten CNN and the ensemble map are available at Neurovault (Gorgolewski et al., 2015; https://neurovault.org/collections/5552/).

*Validating the population-based inference scheme*

#### 2.10.1. Replicability of the produced explanation map as a function of sample size

To examine whether creating explanation maps based on a larger population would increase the split-sample similarity, two population-level explanation maps were created by sampling *m* subjects (*m* = 1, 6, 11,…,101) with replacement from two groups. The groups were created by randomly splitting to half, a sample of 200 subjects from the test and validation sets. Each map was thresholded, binarized (see 2.8) and the Dice similarity and the MHD were calculated between the two maps as a function of the sample size *m*. The procedure was repeated 100 times, and for each iteration, the 200 subjects were randomly assigned to the two groups. This test was repeated independently for each population-based explanation map derived from the ten CNNs.

#### 2.10.2. Benchmarking the results against a standard voxel-based morphometric analysis

To examine whether the derived explanation map elicits similar regions to those detected with established methods, we compared it with a baseline obtained from studies that use voxel-based morphometry (VBM; Ashburner & Friston, 2000) to test structural age-related changes. Briefly, in the VBM method, a mass-univariate test between the tissue composition of any voxel in the brain and a given external variable (age) is conducted. To address the differences in brain position and anatomy, all brain volumes are normalized to a common space and then smoothed by a Gaussian kernel to account for small registration differences. In the current study, we used a published activation likelihood estimation (ALE) meta-analysis of age VBM studies (Vanasse et al., 2018). Here, by utilizing peak reported coordinates from several VBM studies (n = 43), the ALE analysis assigns each voxel the probability that it lies within a reported peak (Laird, Bzdok, Kurth, Fox, & Eickhoff, 2011). The ALE value across all the superthreshold (1^st^ percentile) voxels in the ensemble population-based explanation map were averaged. This empirical value was compared to a null distribution created by randomly sampling one percent of the voxels within the brain mask.

#### 2.10.3. Specificity of the regions obtained in the analysis to the employment of the current model

To evaluate the contribution of the regions discovered using the population-based explanation map to the prediction of the current model we examined how variability in their age-controlled volume correlated with the model’s prediction error. Regional volumes of cortical and subcortical areas were extracted using the Desikan-Killiany atlas (Desikan et al., 2006) computed using Freesurfer following by regressing out the total intracranial volume (ICV; Voevodskaya et al., 2014). Next, subjects’ chronological age was further regressed out from these values to produce the age-normalized volume. Prediction error was formulated as the signed difference between the chronological age and the predicted age. The test was conducted separately for each anatomical ROI in the parcellation. Specific ROIs within the Desikan-Killiany parcellation, such as WM hyperintensities and the fifth ventricle, exist for only some of the subjects (n= 619; from the test/validation sets), and thus were subsequently excluded (9 excluded, 98 remained; see supplementary Fig. 6 for the complete list).

## 3. Results

We start by presenting the model’s ensemble performance for predicting subject chronological age from their T1 structural images on an unseen test set (N = 526). Then, using a novel inference scheme, we locate the anatomical regions that contributed the most to the model’s prediction. We validate the robustness of our inference framework in three ways. First, we demonstrate that it substantially increases reliability compared to previous methods, creating more coherent and localized explanation maps. Second, we quantitatively compare these explanation maps to age voxel-based morphometric studies, demonstrating significant overlap with a simple baseline model. Finally, we demonstrate that this approach enables to gain specific insights about the model by identifying brain regions for which the model shows the highest sensitivity to inter-subject volumetric variability.

### 3.1. Estimating “Brain Age”

Several attempts were previously made to identify the relation between chronological age and brain structure, using various feature extraction techniques, advanced preprocessing methods and a relatively limited sample size (Irimia, Torgerson, Goh, & Van Horn, 2015; Kandel, Wolk, Gee, & Avants, 2013; Shamir & Long, 2016). Here, we build upon recent progress in utilizing CNNs for predicting chronological age from raw structural brain imaging (Cole & Franke, 2017) and introduce substantial improvements using an ensemble of models. In the current work, ten randomly initialized CNNs, were separately trained. The mean MAE across networks was 3.72 years (± 0.17), and the Pearson correlation between the predicted and the chronological age was 0.97 (±0.001). Next, a simple linear regression model was trained on the output of each network to find an optimal linear combination between them, yielding an MAE of 3.07 and a Pearson correlation of 0.98 to the chronological age (Fig. 3; see supplementary Fig. 1 for evaluation per data set).

**Figure 3.**
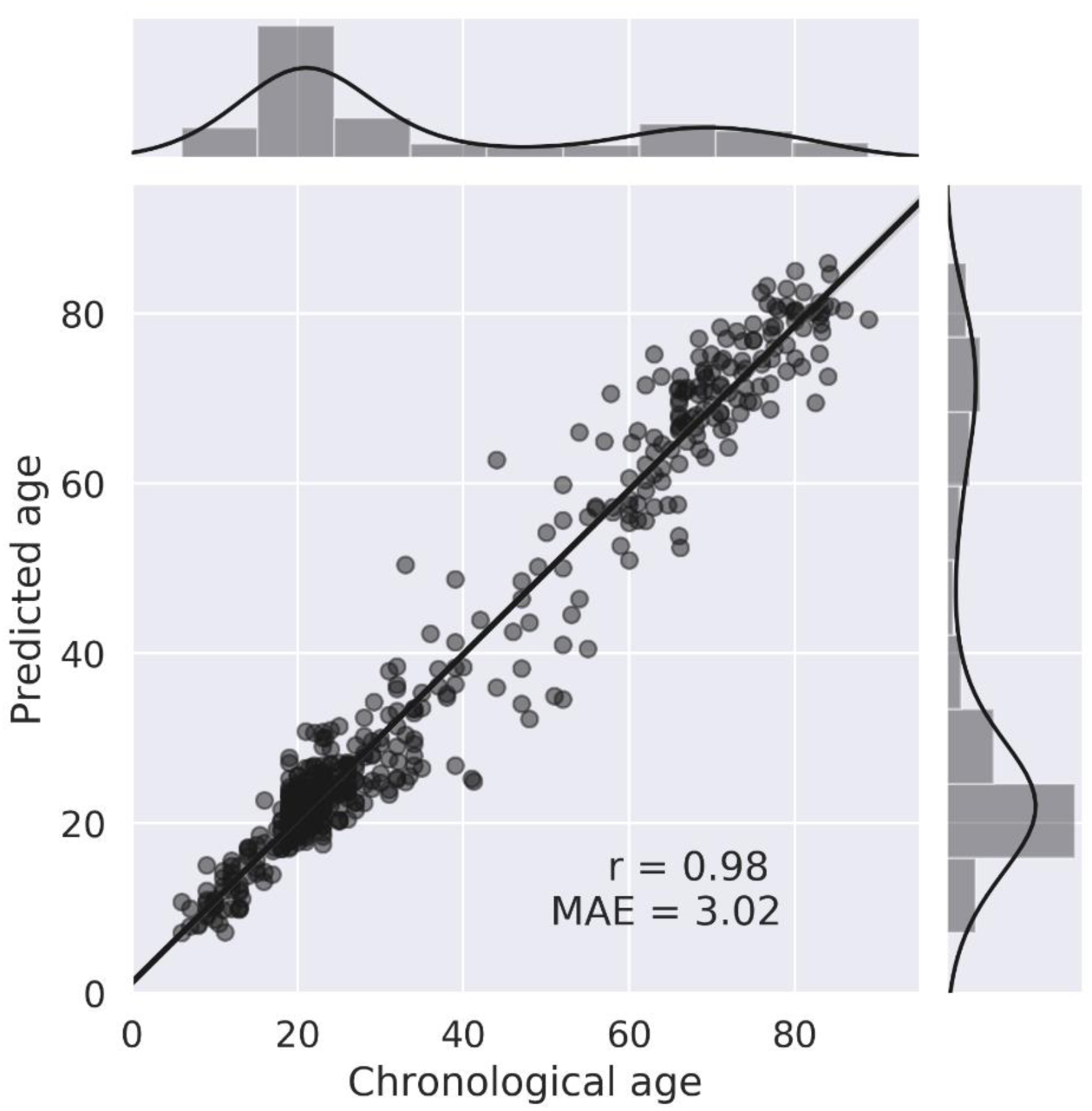
Regression plot of the chronological age compared to the model’s prediction for the test set. The main plot depicts the Pearson correlation coefficient between the chronological and the predicted age; the Pearson correlation coefficient (r) and the mean absolute error (MSE) are indicated on the plot. The data points are presented with partial transparency thus overlapping points are shown in darker gray. The top and right panels of the figure depict histograms and kernel density plots of the distribution of the chronological age and the predicted age (respectively) obtained in the test set.

### 3.2. From the model to the brain – a novel inference scheme

Building upon previous attempts to assign pixel-wise (or voxel-wise) explanation measures to a model’s prediction (Smilkov et al., 2017), we propose that creating explanation maps based on a population, rather than on a specific sample, may substantially improve the coherence and reliability of these maps. We create these maps for each of the ten independently trained CNNs and examine their similarity. Then, using an aggregated map across all ten networks we present the brain regions that contributed the most to predicting age.

#### 3.2.1. Assessing the similarity of explanation maps within the ensemble

Explanation maps for the ten CNNs, averaged across 100 subjects from the test/validation set were produced to create a population-based map (see 2.7; supplementary Fig. 2). First, to assess the similarity between each pair of these produced maps among the ten independently trained networks, each map was thresholded (5^th^ percentile) and binarized, then the Dice coefficient similarity measure was computed for each pair of maps (supplementary Fig. 3). We found a significant Dice similarity across all 45 possible pairs (Dice coefficient: m= 0.17, SD =0.058; binomial test: p < 0.001). Since Dice similarity fails to capture the relation between two maps that are adjacent in the Euclidean space but non-overlapping, we additionally computed the MHD (Dubuisson & Jain, 2002), taking the symmetric average surface distance, among all pairs. We found that the mean MHD among all possible pairs was 6.44 mm (SD = 1.22). Thus, even though these different population-based maps were derived from independently trained networks, there is a moderate, significant, overlap between them. The fact that this overlap is merely moderate coincide with the prediction differences that allow the accuracy gain in ensemble prediction.

#### 3.2.2. Mapping the anatomical regions underlying “brain age” prediction

After estimating the similarity among the different explanation maps for the ten CNNs, we created an ensemble population-based map by taking the median value for each voxel across all networks. We report how the ES is distributed among different tissue types, and among different anatomical regions in order to examine their contribution for age prediction. Testing the volume-normalized contribution of each tissue type, we found that cavities containing CSF and choroid plexus had the highest contribution (35.62%), followed by subcortical GM (27.66%), WM (19.49%), and finally, cortical GM (17.23%) which contributed the least. Table 2 presents the location of clusters (> 100 voxels) in the threshold explanation map (1^st^ percentile). We found that the structures contributing most to age prediction in our model were the ventricles, subarachnoid cisterns, and their borders (see Fig. 4). Specifically, the 4^th^ ventricle, the ambient cistern bilateral to the midbrain, the superior cerebellar cistern, the bilateral Sylvian cistern, the lateral ventricles, the interpeduncular cistern, and the right parahippocampal fissure. WM tracks that were found important for age prediction were the bilateral tapetum, the right anterior limb of the internal capsule and the left medial lemniscus. Finally, the bilateral thalamus and the right precentral gyrus were the GM regions that contributed most to the prediction. Both these analyses support the notion that age prediction in the current model is largely based on age-related morphological changes in the cavities containing CSF.

**Figure 4.**
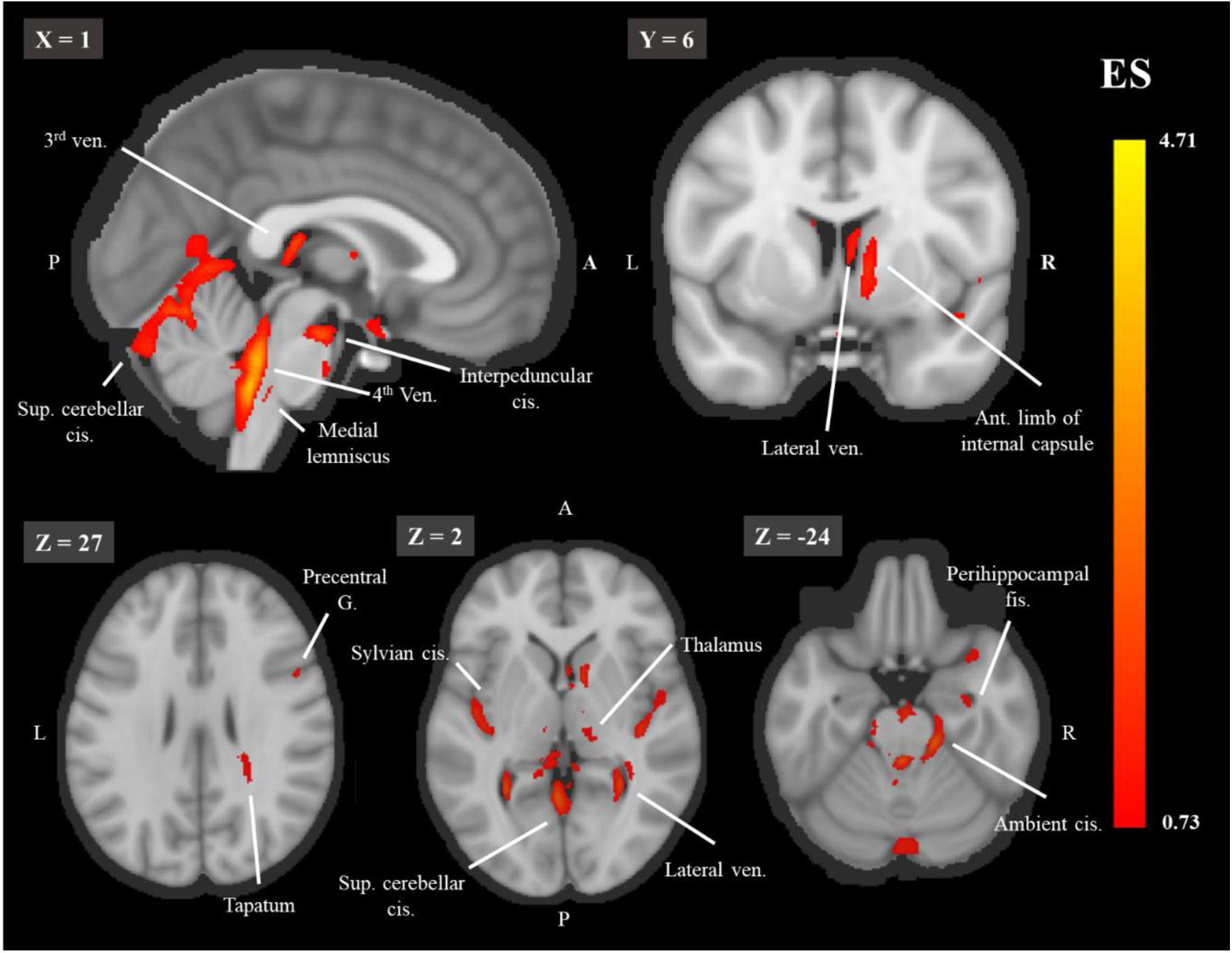
The threshold explanation map shown on a midsagittal (top left), a coronal (top row left) and 3 axial (bottom row) slices. Aggregated explanation map across 100 subjects and the ten networks, thresholded for the 1^st^ percentile of the ES. Abbreviations: ant. = anterior, cis. = cistern, g. = gyrus, fis. = fissure, ven. = ventricle. For each image, the slice number in the MNI template is indicated on the left upper corner. The color bar indicates the values of the ES.

**Table 2.** Anatomical location of clusters in the threshold explanation map. MNI coordinates of clusters in the ensemble population-based explanation map threshold for the 1^st^ percentile. Cluster size is reported in terms of the number of voxels, and the peak ES is the maximum ES within the cluster.

**MNI coordinates**
| **Region** | **Cluster size** | **x** | **y** | **z** | **Peak ES** |
| --- | --- | --- | --- | --- | --- |
| 4th ventricle and ambient cistern | 5531 | -2 | -43 | -39 | 4.71 |
| Superior cerebellar cistern | 4619 | -3 | -55 | 0 | 2.54 |
| R tapetum | 1984 | 27 | -43 | 18 | 1.35 |
| L Sylvian cistern | 1409 | -45 | -16 | 11 | 1.62 |
| R Lateral ventricle | 1115 | 6 | 1 | 8 | 1.42 |
| R Sylvian cistern | 1105 | 41 | -20 | 0 | 1.56 |
| R Lateral ventricle | 1024 | 30 | -49 | 2 | 1.83 |
| R Anterior limb of internal capsule | 847 | 11 | 6 | 2 | 1.48 |
| 3rd ventricle | 787 | 0 | -26 | 11 | 1.89 |
| L lateral ventricle | 740 | -28 | -52 | 4 | 2.08 |
| Interpeduncular cistern | 446 | 1 | -17 | -22 | 2.09 |
| R Sylvian cistern | 440 | 39 | 12 | -20 | 1.23 |
| L medial lemniscus | 307 | -2 | -36 | -41 | 1.25 |
| L thalamus | 303 | -12 | -18 | 11 | 0.993 |
| L ambient cistern | 238 | -12 | -34 | -13 | 1.18 |
| Tapatum left | 231 | -26 | -49 | 15 | 0.989 |
| R thalamus | 212 | 13 | -17 | -2 | 0.925 |
| L ambient cistern | 171 | -16 | -24 | -21 | 1.17 |
| R precentral gyrus | 127 | 50 | 11 | 29 | 0.958 |
| Perihippocampal fissure | 110 | 32 | -9 | -26 | 1.06 |

### 3.3. Validating the population-based inference scheme

To validate the suggested approach for detecting regional contribution to a CNNs ensemble, we conducted three tests. First, we tested the importance of sample size to the explanation maps replicability by testing the half-split similarity of these maps as a function of the population size. Second, to test whether these results coincide with data from other studies, we tested the similarity of the produced maps to a simple baseline of VBM studies meta-analysis. Lastly, to confirm the specificity of these results to the current model, we examined whether the produced maps highlight the particular brain regions for which the model shows the highest sensitivity to inter-subject volumetric variability.

#### 3.3.1. Replicability of the produced explanation map as a function of sample size

It is not clear to what extent an explanation map derived from a single sample would indeed represent the entire population. To examine this issue, we tested the split-sample similarity of the explanation maps with a gradually increasing sample size obtained from two separate groups. In each test repetition (k = 100), the groups were created by randomly half-splitting a sample of 200 subjects (see 2.10). We report the Dice similarity and the MHD among these maps as a function of the sample size drawn from them (see Fig. 3). Across all ten networks, we found an increase of the Dice similarity and a decrease of the MHD as a function of the sample size, ranging from a single sample to 101 samples (mean Dice = 0.19, mean MHD = 3.73; mean Dice = 0.74, mean MHD = 1.00; respectively) (Fig. 5a, 5b). The relative improvement in the replicability of these maps asymptotes at 40-60 subjects, such that adding more samples had little further impact. Figure 5c shows 2D glass brain projections of the population-based maps to illustrate the change as a function of the sample size, resulting in a visually apparent increase in coherence and decrease in noise. These results suggest that whether due to noise or fundamental differences in subject-specific maps, such as gender or age group (Jäncke et al., 2015; Tamnes et al., 2013), an explanation produced from a single sample has low replicability. Thus, when addressing a general, rather than a sample-specific question, a population-based explanation should be favored, as opposed to previous practices in the field (Zhang & Zhu, 2018).

**Figure 5.**
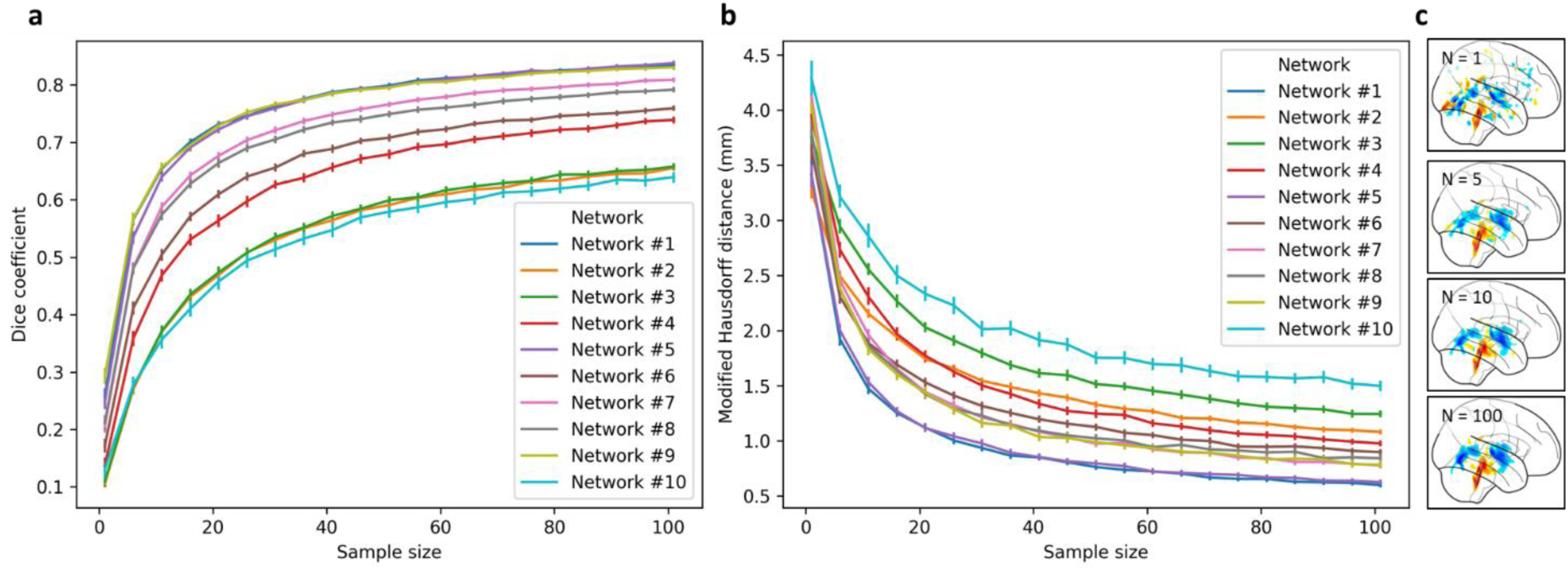
The split-sample similarity of the explanation maps as a function of sample size. The similarity of two maps produced from an increased sample size from two separate groups (n=100 for each group) was measured using (a) Dice coefficient and (b) MHD (mm). The results are reported for all ten CNNs and each is presented in a separate color. (C) A visual illustration of an explanation map for network 1 produced by increasing the sample size (from top to bottom, N = 1,5,10,100). The error bars represent a 95% confidence interval.

**Figure 6.**
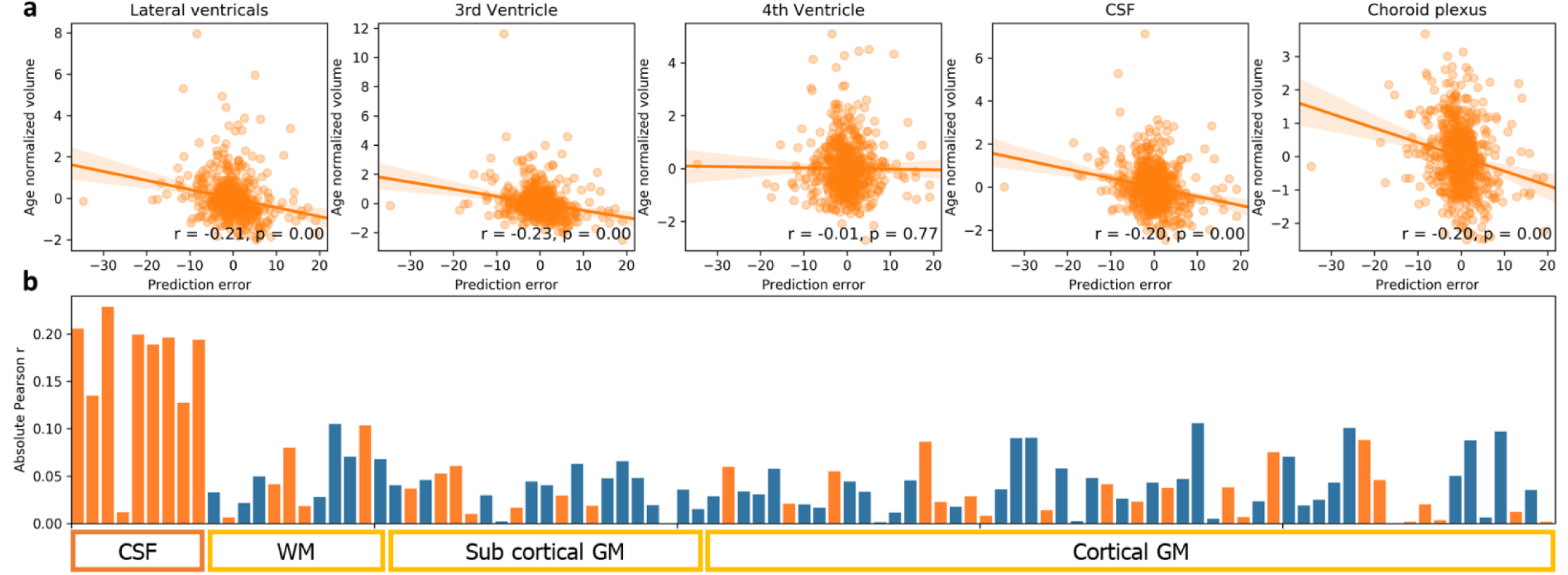
Deviation in volume from age norm and prediction error. (a) Graphs of five ROIs, detected with the current inference scheme, showing the correlation between the age-controlled volume and the signed prediction error. Age-normalized volume was computed by regressing out subjects chronological age from the measured volume. Volume was determined according to the Desikan-Killiany atlas fitted with Freesurfer. Prediction error was formulated as the chronological age minus the predicted age. Note that for the sake of brevity, in the upper five plots, the volume of the lateral ventricles and choroid plexus was computed as the sum of their sub-parcellations. (b) The bar graph depicts the correlation between the age normalized volume and the signed prediction error for all the 98 regions in the parcellation. Positive correlations are presented in blue and negative in orange for simple magnitude comparison. As shown, the age-controlled volume of cavities containing CSF and the choroid plexus (L/R Lateral Ventricle, L/R inferior Lateral Ventricle, 3rd ventricle, non-ventricles CSF, L/R choroid plexus), except for the 4^th^ ventricle, had the largest correlation with the model’s prediction error compared with all other WM/GM regions (see supplementary for a figure with the full labels).

#### 3.3.2. Benchmarking the results against a standard voxel-based morphometric analysis

To quantitatively assess whether the regions detected in the current inference scheme coincide with previous findings, we compared the resulting explanation maps to a baseline obtained from VBM studies testing structural age-related changes. The VBM method has the desired property of allowing to test the relation between the estimated tissue composition of each voxel and any given relevant variable, age in the present case. In contrast, other methods are limited to a specific set of ROIs or a given brain parcellation that often fails to properly parcellate non-GM regions. We used a published ALE meta-analysis of age VBM studies (n = 15, Vanasse et al., 2018) in which each voxel is assigned with a probability for its location in a reported peak coordinate in one of the studies (Laird et al., 2011). Using this map, we examined whether that ALE value is significantly higher within the regions identified using the threshold explanation map with a permutation test. The mean ALE value within the super-threshold (1^st^ percentile) explanation map was higher than any set of randomly selected voxels in the permutation test (k = 10,000; meta-analysis: empirical mean ALE: 0.003, p<0.0001; see supplementary, Fig. 4). Interestingly, both methods highlighted regions surrounding the lateral and 3^rd^ ventricles, subcortical areas, and the bilateral insulas/sylvian cisterns, as opposed to cortical regions that appeared only in the ALE map (see supplementary, Fig. 5).

#### 3.3.3. Specificity of the regions obtained in the analysis to the employment of the current model

Prediction errors could result from the inability of the model to capture the complexity of the brain aging process or due to the natural variability in brain morphology within the population. Exploiting the latter, in the current analysis we aimed to examine whether prediction error could be associated with volumetric variability of specific brain regions. Specifically, by applying the Desikan-Killiany atlas using Freesurfer we tested whether age-controlled volume of the ventricles and cisterns that were highlighted by the inference scheme were correlated with the CNNs ensemble prediction error. Indeed, we found a significant correlation between the age-normalized volume and the prediction error for the ventricles excluding the 4^th^ ventricle, the choroid plexus and non-ventricular CSF (n = 619; for all 9 regions but 4^th^ ventricle: r > 0.13, p < 0.002). This correlation was higher in these regions than in any other brain region in the parcellation supporting the specificity of the results to the regions obtained using the population-based explanation maps (see Fig. 4). Interestingly, this specificity was not apparent when examining the correlation between regional volume and chronological age, in which significant correlation is seen in almost all regions (> 93% of the ROI; see supplementary, Fig. 7). Put differently, while volume of almost all regions correlated with age, deviance from the age norm in the ventricles and CSF, detected using the population-based maps, best reflected the prediction error. This suggests that the current inference scheme not only detected regions that are altered in aging, but it detected the distinct regions that had the highest contribution to the current prediction, attesting to the high specificity of this method.

## 4. Discussion

In the current study, we examined whether individuals’ chronological age could be predicted from T1 MRI scan and whether it is possible to localize the underlying brain regions that allow such prediction. Using a large aggregated sample of 10,176 subjects we trained and validated an ensemble of 3D CNN models, and showed that “brain age” could be estimated from raw T1 MRI with MAE of ∼3.1 years. We demonstrated that the use of an ensemble of models rather than a single estimator reduces the MAE in more than 6 months and provided evidence that such gain is due to the difference in the features or brain regions that are utilized by each model. Brain age was previously shown to be indicative of neurodegenerative diseases and other clinical conditions (Cole & Franke, 2017), thus improving the estimation and the interpretability of this biomarker could be an important step toward integrating it in clinical use.

### 4.1. Identifying the brain regions underlying age prediction using population-based explanation maps

Drawing from previous studies aiming at identifying regional contributions to the model’s prediction, we aimed to locate the brain regions that governed our brain age estimation. Here we presented a novel approach to aggregate multiple explanation maps from several subjects, thus creating a population-based map. This was achieved by deriving a series of transformations warping the 3D volumes presented to the CNN into the MNI space. We then applied those transformations to the computed explanation maps, thus allowing to average different explanation maps in a common space. This approach precludes the need for pre-registration to a common template in the training stage, as done previously (Ceschin et al., 2018), a step that is error-prone, time-consuming and might result in the loss of relevant structural information (Iscan et al., 2015). Importantly, this approach could be used for any ML application for neuroimaging.

In order to validate our method, we tested how it affects three important aspects. First, we quantitively assessed how sample size in population-based maps improved their reproducibility. We reported a substantial improvement in split-sample similarity as moving for a map based on a single subject to a map based on a population of 40-60 subjects. The low split-sample similarity of single-subject maps emphasized the need to apply such classic neuroscience practices when analyzing these explanation maps. Next, we demonstrated that despite the methodological differences, the proposed map shows significant similarity with ALE maps from age VBM meta-analysis study (Vanasse et al., 2018), attesting to its convergence validity. Finally, using regional volumetric measures we demonstrated that brain regions highlighted by our method were found as the ones with the highest influence on the model’s prediction, indicating the specificity of the derived maps to the current model.

### 4.2. Reducing noise or averaging over true relevant population differences?

Comparing our approach for deriving population-based explanation maps to one based on a single sample as in Smilkov et al. (2017) paper, we demonstrate an increase in reproducibility and a distinct visual improvement in these maps’ coherence. We therefore discuss possible mechanisms that may account for these findings. In their study, Smilkov et al. (2017) demonstrated that the derivative of CNNs are highly noisy, and averaging explanation maps derived from several noised samples of the same input can improve these maps. Here, after applying the Smilkov et al. (2017) method, we further averaged multiple explanation maps derived from different inputs, i.e. brain volumes of different subjects. A possible explanation to the apparent reproducibility improvement is that sampling from the true input distribution (brain volumes of different individuals), rather than mere noised samples of the same input, would result in estimation that is more robust to local gradient noise. A second, non-exclusive possible account might stem from the fact that the model was trained on brain volumes from heterogenic population. Differences in brain aging trajectory were found both at the individual level (Raz, Ghisletta, Rodrigue, Kennedy, & Lindenberger, 2010) and among different populations, for example in relation to gender (Jäncke et al., 2015). Thus, it is possible that the model extracts different features due to relevant structural variability in different populations.

### 4.3. The ventricles and cisterns as biomarkers for brain aging

Aging is accompanied by multiple processes affecting the human brain, manifested in structural changes that could in part be quantified by neuroimaging (see: Lorio et al., 2016). Accordingly, a wealth of literature reported a complex pattern of morphological changes evident across all brain regions, but arguably more apparent in some areas such as the frontal lobes, insular cortices, and the hippocampus (Fjell et al., 2009). Interestingly, in our model, it is the ventricles and cisterns that were highlighted as the most relevant for age prediction. Several possible reasons might account for this finding. First, CSF volume was found to increase already from young adulthood (Courchesne et al., 2000), thus it may constitute an early aging biomarker. Notably, since CSF pressure remains relatively constant and even decreases in old age (Fleischman et al., 2012), it is likely that CSF expansion reflects a decrease in WM/GM volumes rather than an increase in CSF pressure. Thus, CSF volume changes might be a surrogate for general brain atrophy, as suggested in previous work (De Vis et al., 2016). Nevertheless, the CNN operation is not likely to be reduced to mere volume extraction of these regions, given the substantially lower accuracy of prediction models based on regional volumetric measurements (Liem et al., 2016; Valizadeh, Hänggi, Mérillat, & Jäncke, 2017). Finally, it is also possible that compared to other brain regions, for example within the cortex, the ventricles and cisterns present lower inter-subject structural variability, thus comprising a more reliable measure of brain aging. These possible reasons could be tested in future work but nevertheless, the ability to generate new biologically relevant hypotheses from a deep learning predictive model is a desirable practice supported here by our novel inference scheme.

### 4.4. Ensemble diversity and similarity among models’ population-based explanation maps

Evidence suggests that prediction based on a set of learning algorithms instead of a single algorithm will result in an accuracy gain (Sagi & Rokach, 2018) that increases as these models are more accurate and diverse (Breiman, 2001; Kuncheva & Whitaker, 2003). Learning diverse models could be achieved by changes in architecture (Singh, Hoiem, & Forsyth, 2016) or introducing different subsets of the training data to each model (Benou, Veksler, Friedman, & Riklin Raviv, 2017). In the context of deep CNN, as opposed to convex or shallow learning algorithms, it has been shown that models that differ only in their random weight initializations constitute an ensemble that is not only adequately diverse, but perform better than models exposed to different subsets of the data (Lakshminarayanan et al., 2017; Lee et al., 2015). In the current work, we examined the similarity among pairs of population-based explanation maps derived from different models within such an ensemble. Although within each model population maps showed high reliability, on average, pairs of models exhibit only moderate similarity. This supports the notion that random weight initializations generate diverse models that utilized different parts of the input, i.e. different brain regions, which may explain the improvement in prediction accuracy. Nevertheless, aggregating these maps revealed that a set of regions, such as the ambient and cerebellar cisterns, consistently utilized across all models. Overall, it seems that general conclusions regarding the contribution of different brain regions to age prediction should be made based on maps derived from multiple models.

### 4.5. Future direction: testing population differences in explanation maps

Computing population-based explanation maps allow examining group differences in maps produced from different populations. For example, one might ask whether a CNN model would extract different aging biomarkers for men versus women or healthy elderly versus individuals diagnosed with AD. These tests could be applied on maps derived from two identical models separately trained on different populations or within the same model trained on both populations. In the second case, subjects’ group affiliation could be explicitly introduced to the model as an input. Alternatively, it will be possible to test whether a distinction among populations in the form of explanation maps differences, would arise without introducing such an input. Importantly, explanation maps yield from a population of subjects, registered to the same template could allow harnessing known neuroscience statistical procedures based on voxel or regional wise comparison of within compared to between-group variability.

### 4.6. Conclusions

Incorporating deep learning for analysis of neuroimaging data requires improvement in both the accuracy of these predictive models and the ability to interpret them, as we aimed to address in the context of age prediction. Respectively, in the current work, we demonstrated that an individual’s chronological age could be estimated with an MAE of 3.1 years from their raw T1 images, yielding a robust biomarker across several datasets. We further showed that aggregating multiple explanation maps substantially increases their reproducibility and allow to create a coherent and localized map depicting and quantifying the contribution of different brain regions to age prediction. From these maps, we conclude that the ventricles and cisterns govern these predictions. We argue that this ability to pinpoint specific brain areas is a key step for utilizing these models as possible brain health biomarkers.

## 5. Data availability

All the data sets used for the model’s training, validation and testing were acquired from open-access data sharing projects. The results of the main analysis, the unthresholded population-based maps for each of the ten CNN and the ensemble map are available at Neurovault (Gorgolewski et al., 2015; https://neurovault.org/collections/5552/).

## 6. Acknowledgments

This research was supported by an Internal funding grant for interdisciplinary research in Ben Gurion University of the Negev to GA and IS.

Our project would not have been possible without several open databases and groups that invested considerable resource and efforts to support neuroimaging data sharing. We wish to acknowledge those study groups and funding agencies:

<u>CORR</u> - Data were provided in part by the Consortium for Reliability and Reproducibility (http://fcon_1000.projects.nitrc.org/indi/CoRR/html/index.html).

<u>ADNI</u> - Data collection and sharing for this project was funded by the Alzheimer’s Disease Neuroimaging Initiative (ADNI) (National Institutes of Health Grant U01 AG024904) and DOD ADNI (Department of Defense award number W81XWH-12-2-0012). ADNI is funded by the National Institute on Aging, the National Institute of Biomedical Imaging and Bioengineering, and through generous contributions from the following: AbbVie, Alzheimer’s Association; Alzheimer’s Drug Discovery Foundation; Araclon Biotech; BioClinica, Inc.; Biogen; Bristol-Myers Squibb Company; CereSpir, Inc.; Cogstate; Eisai Inc.; Elan Pharmaceuticals, Inc.; Eli Lilly and Company; EuroImmun; F. Hoffmann-La Roche Ltd and its affiliated company Genentech, Inc.; Fujirebio; GE Healthcare; IXICO Ltd.;Janssen Alzheimer Immunotherapy Research & Development, LLC.; Johnson & Johnson Pharmaceutical Research & Development LLC.; Lumosity; Lundbeck; Merck & Co., Inc.;Meso Scale Diagnostics, LLC.; NeuroRx Research; Neurotrack Technologies; Novartis Pharmaceuticals Corporation; Pfizer Inc.; Piramal Imaging; Servier; Takeda Pharmaceutical Company; and Transition Therapeutics. The Canadian Institutes of Health Research is providing funds to support ADNI clinical sites in Canada. Private sector contributions are facilitated by the Foundation for the National Institutes of Health (www.fnih.org). The grantee organization is the Northern California Institute for Research and Education, and the study is coordinated by the Alzheimer’s Therapeutic Research Institute at the University of Southern California. ADNI data are disseminated by the Laboratory for Neuro Imaging at the University of Southern California.

<u>GSP</u> - Data were provided in part by the Brain Genomics Superstruct Project of Harvard University and the Massachusetts General Hospital, (Principal Investigators: Randy Buckner, Joshua Roffman, and Jordan Smoller), with support from the Center for Brain Science Neuroinformatics Research Group, the Athinoula A. Martinos Center for Biomedical Imaging, and the Center for Human Genetic Research. 20 individual investigators at Harvard and MGH generously contributed data to GSP.

<u>FCP</u> - Data were provided in part by the Functional Connectomes Project (https://www.nitrc.org/projects/fcon_1000/).

<u>ABIDE</u> - Primary support for the work by Adriana Di Martino, and Michael P. Milham and his team was provided by the NIMH (K23MH087770), the Leon Levy Foundation, Joseph P. Healy and the Stavros Niarchos Foundation to the Child Mind Institute, NIMH award to MPM (R03MH096321), National Institute of Mental Health (NIMH 5R21MH107045), Nathan S. Kline Institute of Psychiatric Research), Phyllis Green and Randolph Cowen to the Child Mind Institute.

<u>PPMI</u> - Data used in the preparation of this article were obtained from the Parkinson’s Progression Markers Initiative (PPMI) database (www.ppmi-info.org/data). For up-to-date information on the study, visit www.ppmi-info.org. PPMI – a public-private partnership – is funded by the Michael J. Fox Foundation for Parkinson’s Research and funding partners, including [list of the full names of all of the PPMI funding partners can be found at www.ppmi-info.org/fundingpartners].

<u>ICBM</u> - Data used in the preparation of this work were obtained from the International Consortium for Brain Mapping (ICBM) database (www.loni.usc.edu/ICBM). The ICBM project (Principal Investigator John Mazziotta, M.D., University of California, Los Angeles) is supported by the National Institute of Biomedical Imaging and BioEngineering. ICBM is the result of efforts of co-investigators from UCLA, Montreal Neurologic Institute, University of Texas at San Antonio, and the Institute of Medicine, Juelich/Heinrich Heine University - Germany.

<u>AIBL</u> - Data used in the preparation of this article was obtained from the Australian Imaging Biomarkers and Lifestyle flagship study of ageing (AIBL) funded by the Commonwealth Scientific and Industrial Research Organisation (CSIRO) which was made available at the ADNI database (www.loni.usc.edu/ADNI). The AIBL researchers contributed data but did not participate in analysis or writing of this report. AIBL researchers are listed at www.aibl.csiro.au.

<u>SLIM</u> - Data were provided by the Southwest University Longitudinal Imaging Multimodal (SLIM) Brain Data Repository (http://fcon_1000.projects.nitrc.org/indi/retro/southwestuni_qiu_index.html).

<u>IXI</u> - Data were provided in part by the IXI database (http://brain-development.org/).

<u>OASIS</u> - OASIS is made available by Dr. Randy Buckner at the Howard Hughes Medical Institute (HHMI) at Harvard University, the Neuroinformatics Research Group (**<u>NRG</u>**) at Washington University School of Medicine, and the Biomedical Informatics Research Network (**<u>BIRN</u>**). Support for the acquisition of this data and for data analysis was provided by NIH grants P50 AG05681, P01 AG03991, P20 MH071616, RR14075, RR 16594, U24 RR21382, the Alzheimer’s Association, the James S. McDonnell Foundation, the Mental Illness and Neuroscience Discovery Institute, and HHMI.

<u>CNP</u> - Data used in the preparation of this article were obtained from the Consortium for Neuropsychiatric Phenomics (NIH Roadmap for Medical Research grants UL1-DE019580, RL1MH083268, RL1MH083269, RL1DA024853, RL1MH083270, RL1LM009833, PL1MH083271, and PL1NS062410). This data was obtained from the OpenfMRI database. Its accession number is ds000030

<u>COBRE</u> - Data were provided by the Center for Biomedical Research Excellence (COBRE) (http://fcon_1000.projects.nitrc.org/indi/retro/cobre.html).

<u>CANDI</u>- Data were provided in part by the Child and Adolescent NeuroDevelopment Initiative (CANDI; Kennedy et al., 2012) - Schizophrenia Bulletin 2008 project.

<u>Brainomics</u> - Data were provided in part by the Brainomics project (http://brainomics.cea.fr/).

<u>NITRC</u> – Data used in this manuscript was partially accessed through the Neuroimaging Informatics Tools and Resources Clearinghouse. NITRC is funded by the NIH Grant numbers: 2R44NS074540 and 1U24EB023398a.

<u>IDA</u> – Data used in this manuscript was partially accessed through the LONI Image and Data Archive. IDA is funded by the NIH and the NIBIB grant numbers P41EB015922 U54EB020406.

