## Supplementary material for "From a deep learning model back to the brain - inferring morphological markers and their relation to aging": Supplamentary materials

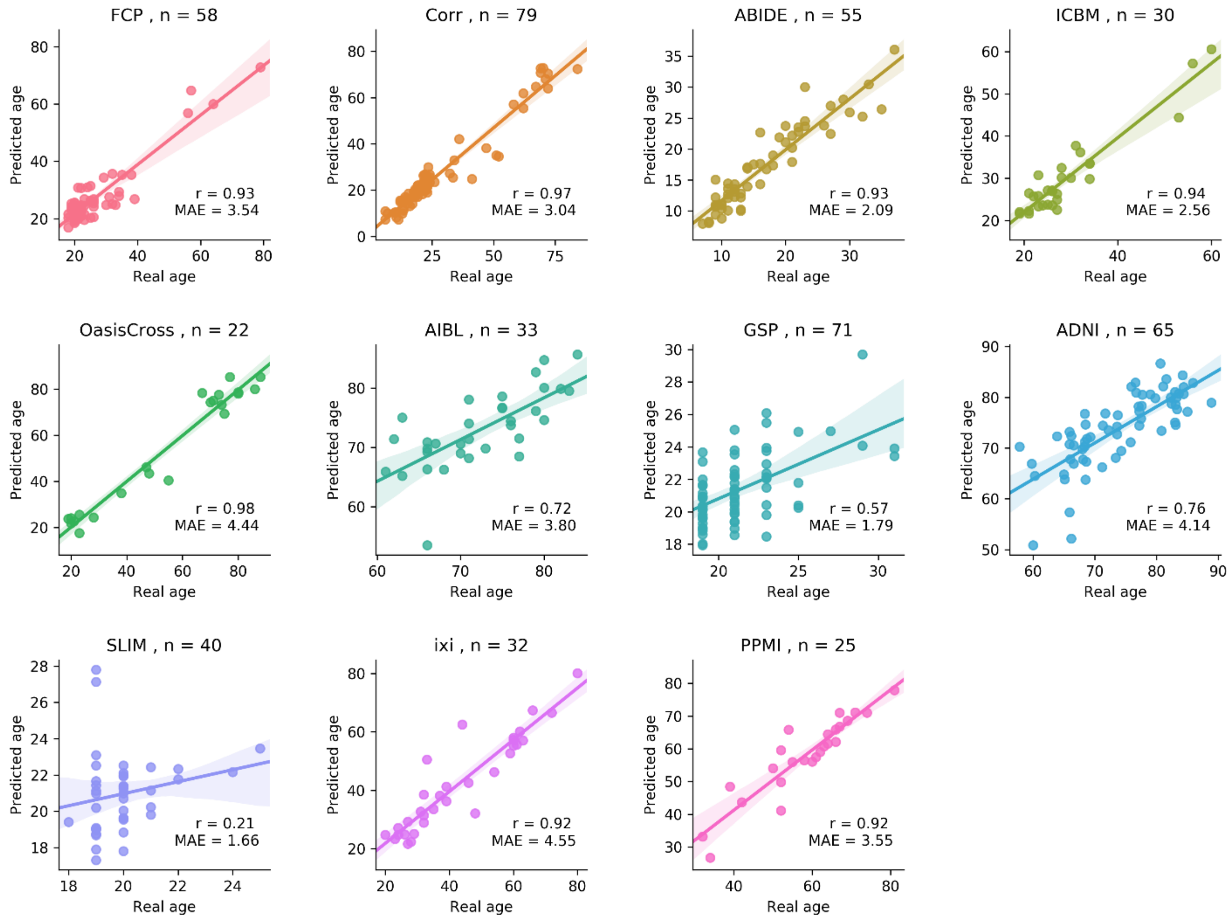


**Figure 1. Regression plot of the chronological age compared to the model's prediction for each one of the large databases employed in the study. The plots depict the Pearson correlation coefficient between the real and the predicted age; r values and mean absolute error (MSE) are indicated on the plot. The title of each plot states the relevant database and number of subjects from the test set. To prevent participants' identification in the GSP study age was rounded to the closest 2 years bin.**

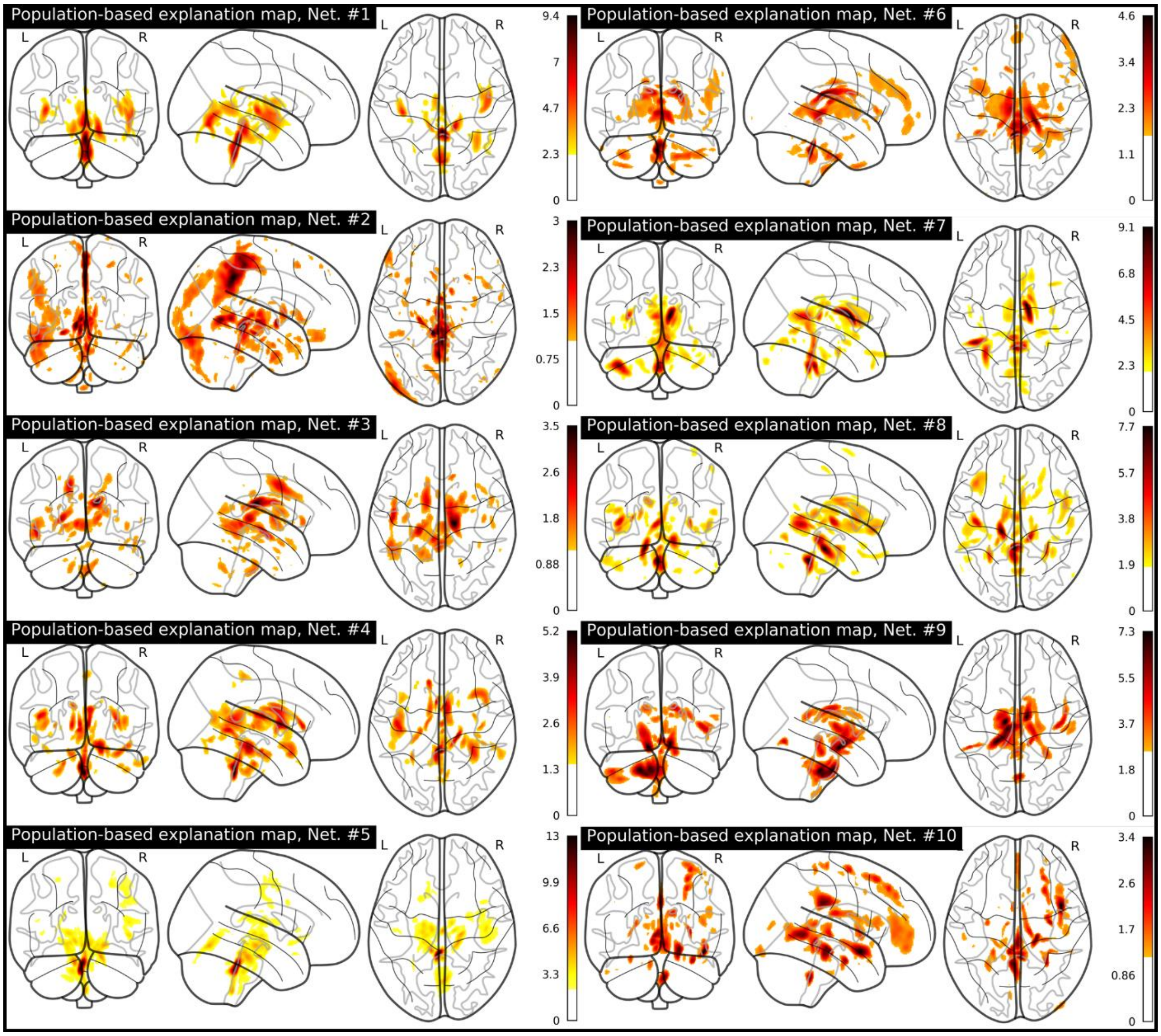


**Figure 2. Coronal (left), sagittal (middle) and horizontal (right) glass brain projections of all population maps derived from the ten networks. Explanation maps were created by averaging 100 subjects-based maps and were threshold to present only 1% of the highest valued voxels within the skull. Colors represent the value of the explanation score (ES).**

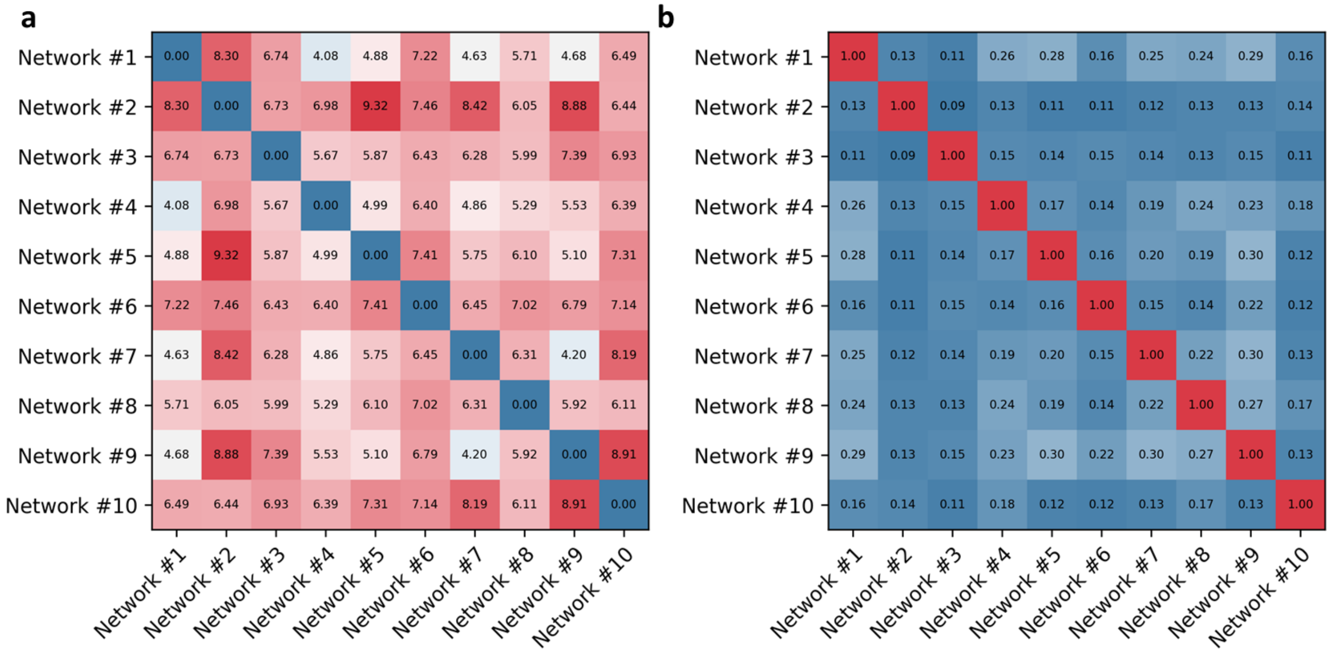


**Figure 3. Distance or similarity among independently trained CNNs for age prediction. (a) Distance matrix of the Modified Hausdorff Distance among all pairs of networks. (b) Similarity matrix of the Dice score among all pairs of networks.**

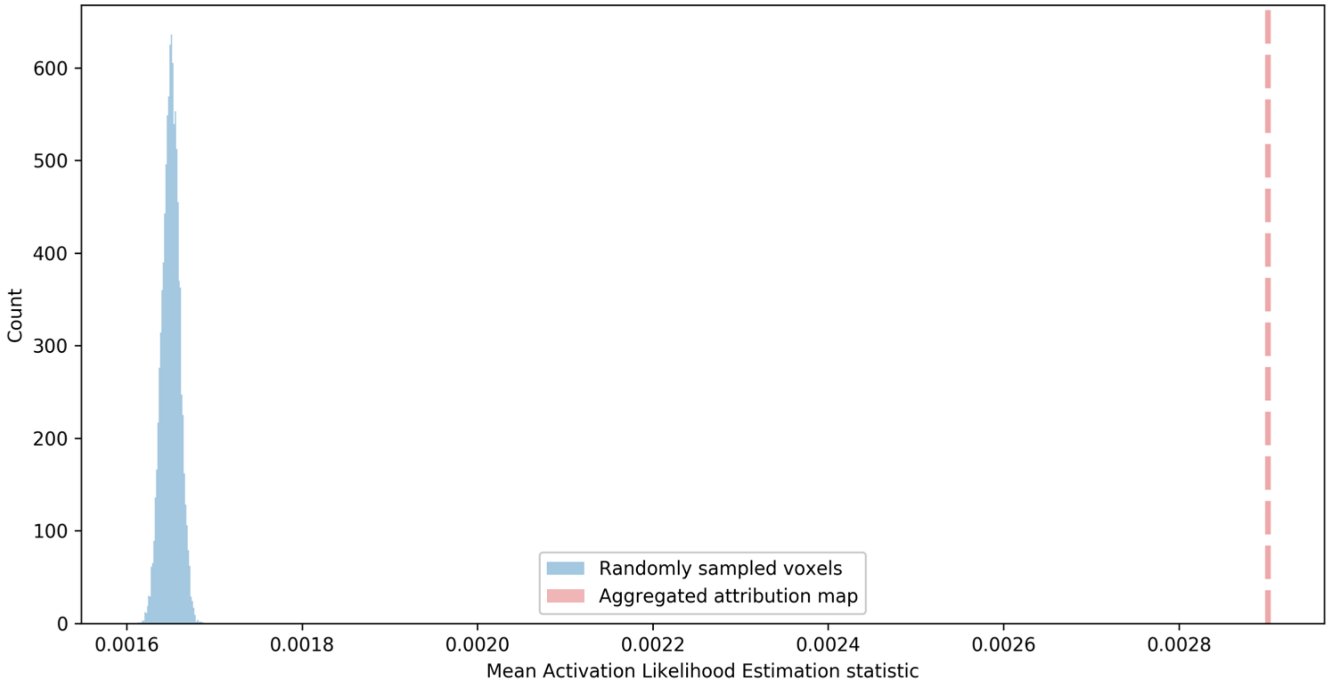


**Figure 4. Histogram of the randomly sampled mean ALE statistics compared to the empirical mean within the super-threshold voxels. In blue a distribution of the mean ALE within the randomly sampled voxels (k =10,000), the red dotted line represents the empirical mean ALE within the super-threshold. All randomly sampled values were smaller than the empirical value.**

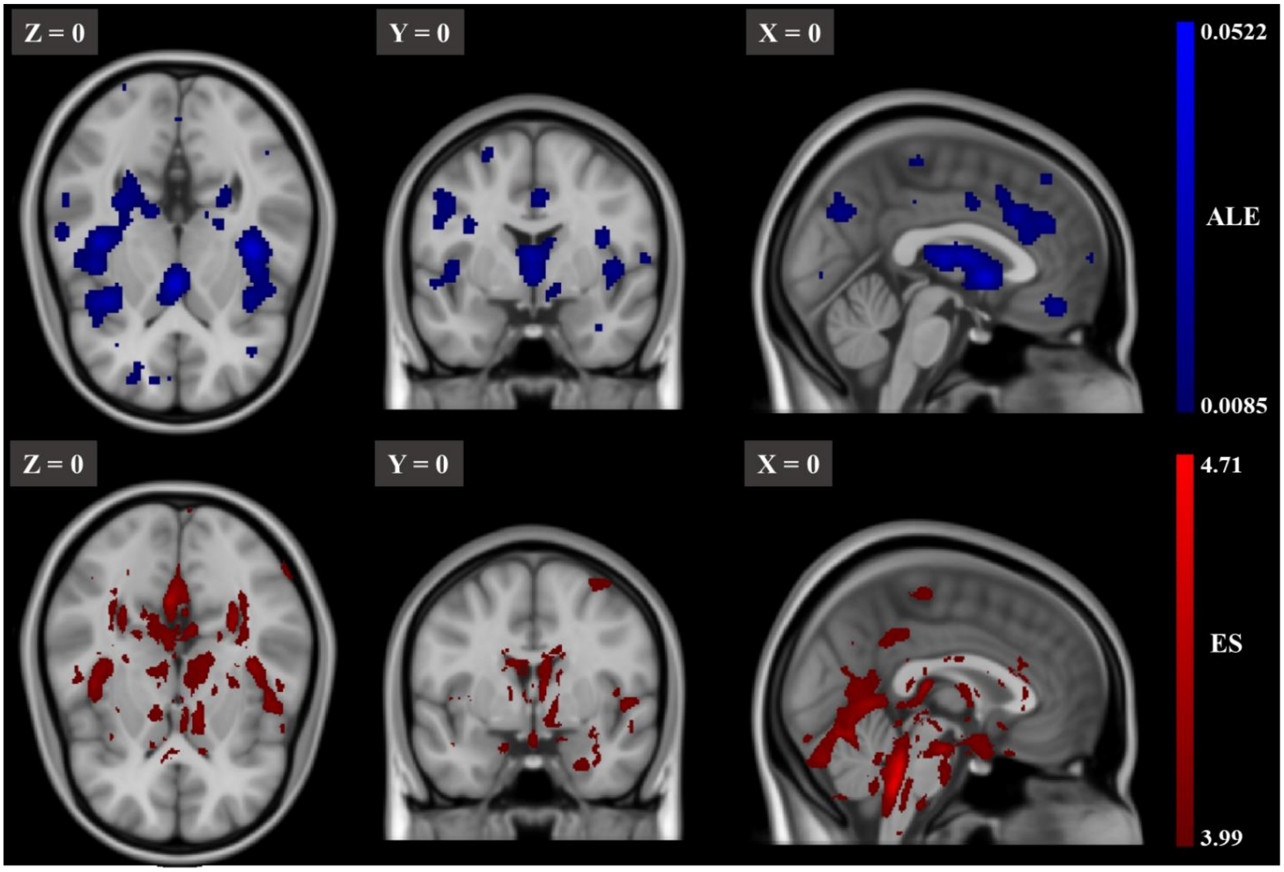


**Figure 5. The threshold "explanation map" and ALE map shown on midsagittal (right), coronal (middle) and axial (left) slices. (top) Threshold ALE map (1st percentile) from Vanasseet al (2018). (bottom) Aggregated, threshold (1st percentile) "explanation map" across 100 subjects and all the 10 networks. For each image, the slice number in the MNI template is indicated on the left upper corner. The color bar indicates the values of the ALE (blue) and the explanation score (ES; red).**

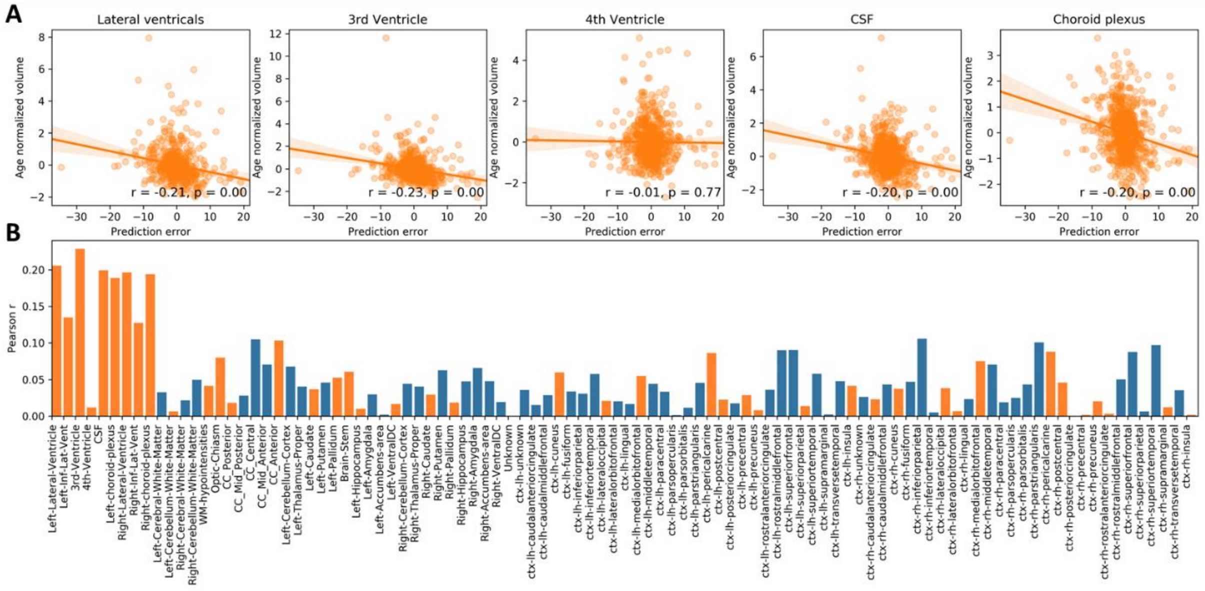


**Figure 6. Deviation in volume from age norm and prediction error. (a) Graphs of five ROIs, detected with the current inference scheme, showing the correlation between the age-controlled volume and the signed prediction error. Age-normalized volume was computed by regressing out subjects chronological age from the measured volume. Volume was determined according to the Desikan-Killiany atlas fitted with Freesurfer. Prediction error was formulated as the chronological age minus the predicted age. Note that for the sake of brevity, volume of the lateral ventricles and choroid plexus was computed as the sum of their sub-parcellations. (b) The bar graph depicts the correlation between the age normalized volume and the signed prediction error for all the 98 regions in the parcellation. Positive correlation is presented in blue and negative in orange for simple magnitude comparison. As shown, the age-controlled volume of cavities containing CSF and the choroid plexus (L/R Lateral Ventricle, L/R inferior Lateral Ventricle, 3rd ventricle, non-ventricles CSF, L/R choroid plexus), except for the 4th ventricle, had the largest correlation with the model's prediction error compared with all other WM/GM regions (see supplementary for a figure with the full labels).**

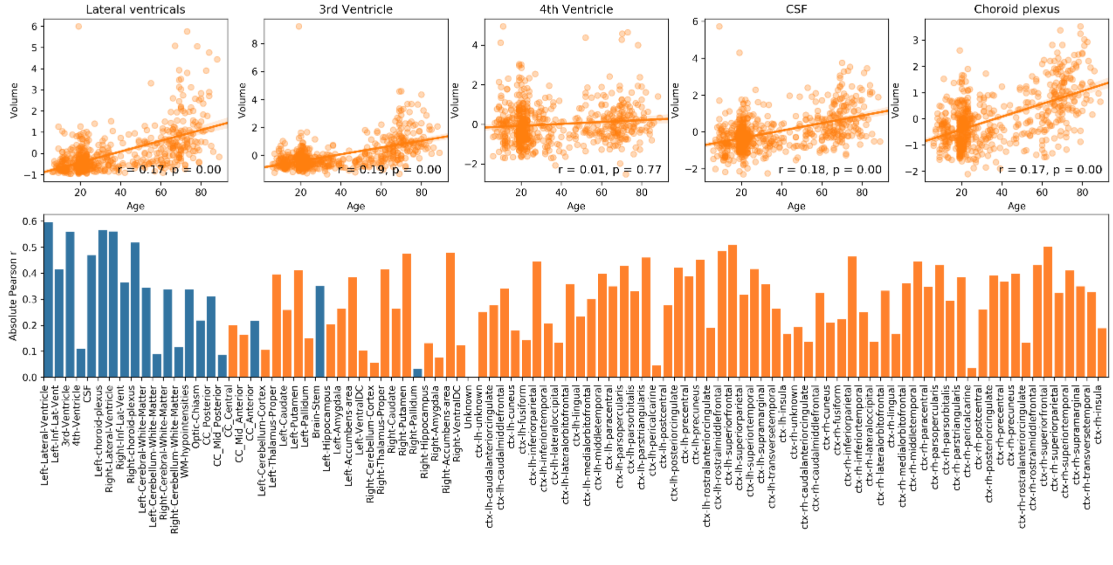


**Figure 7. Correlation of volume and age. (a) Graphs showing the correlation between each ROI volume and chronological age. Volume was determined according to the Desikan-Killiany atlas fitted with Freesurfer (Desikan et al., 2006). . Note that for the sake of brevity, the volume of the lateral ventricles and choroid plexus was computed as the sum of their sub-parcellations. (b) The bar graph depicts the value of the correlation coefficient between volume and age for all the 98 regions in the parcellation. Positive correlation is presented in blue and negative in orange for simple magnitude comparison.**
